# Luigi: Large-scale histopathological image retrieval system using deep texture representations

**DOI:** 10.1101/345785

**Authors:** Daisuke Komura, Keisuke Fukuta, Ken Tominaga, Akihiro Kawabe, Hirotomo Koda, Ryohei Suzuki, Hiroki Konishi, Toshikazu Umezaki, Tatsuya Harada, Shumpei Ishikawa

## Abstract

**Background:** As a large number of digital histopathological images have been accumulated, there is a growing demand of content-based image retrieval (CBIR) in pathology for educational, diagnostic, or research purposes. However, no CBIR systems in digital pathology are publicly available.

**Results:** We developed a web application, the Luigi system, which retrieves similar histopathological images from various cancer cases. Using deep texture representations computed with a pre-trained convolutional neural network as an image feature in conjunction with an approximate nearest neighbor search method, the Luigi system provides fast and accurate results for any type of tissue or cell without the need for further training. In addition, users can easily submit query images of an appropriate scale into the Luigi system and view the retrieved results using our smartphone application. The cases stored in the Luigi database are obtained from The Cancer Genome Atlas with rich clinical, pathological, and molecular information. We tested the Luigi system by querying typical cancerous regions from four cancer types, and confirmed successful retrieval of relevant images.

**Conclusions:** The Luigi system will help students, pathologists, and researchers easily retrieve histopathological images of various cancers similar to those of the query image.

## Background

Recently, computational image analysis has been applied to the ever increasing volume of digital histopathological and medical images for many purposes[1,2]. Content-based image retrieval (CBIR) is one such application that facilitates retrieval of the most relevant images to a query image from an image database. In digital pathology, CBIR systems are useful in many situations, particularly diagnosis, education, and research. For example, CBIR systems can be used for educational purposes by students and beginner pathologists to retrieve relevant cases or histopathological images of tissues. In addition, such systems are also helpful to professional pathologists, particularly with the diagnosis of rare cases.

Several CBIR systems to retrieve similar histopathological images have been proposed[3,4]; however, to the best of our knowledge, no publicly available system has yet been released. Therefore, we developed Luigi, a novel web-based CBIR system, which allows users to easily retrieve similar histopathological images of cancer cases with various useful annotations. The Luigi system calculates image relevance based on representations from a deep convolutional neural network (CNN), which has achieved great success in a wide range of visual tasks. Well-trained deep neural networks provide visual features that perform well in a wide range of computer vision applications, including medical image analysis[5–8]. Although some CBIR applications have attempted to employ deep learning, such applications require label information, which is difficult to obtain in the field of pathology and is not scalable to large and various datasets. Based on the observation that pathological images, particularly of various cancer cases, potentially have textural structures and that spatial autocorrelation is often used in texture analysis, rather than using features from a CNN directly, we adopted the autocorrelation of the feature maps of a CNN pre-trained on ImageNet, which is referred to as deep texture representations[9]. This approach works remarkably well with pathological images without the necessity of additional learning processes for specific datasets. Therefore, the Luigi system is easily applicable to various tissues and scalable to a large number of histopathological images. A smartphone application was also developed that allows users to easily submit cases and view the retrieved results without the need for a dedicated system. Moreover, as all cancer cases in Luigi are retrieved from The Cancer Genome Atlas (TCGA)[10], which is a collection of more than 10000 whole slide images (WSIs) of 32 cancer types with rich clinical, pathological, and molecular information, pathologists and researchers can also obtain further insight into the molecular basis of the cancer of interest. The effectiveness of the Luigi CBIR system was demonstrated by querying typical cancerous regions.

## Implementation

The Luigi system was developed with a Python back end and Clojure front end; thus, it can be accessed by any modern web browser with a Mac OS X, or Windows operating system. An overview of the Luigi system is shown in Figure 1.

**Figure 1.**
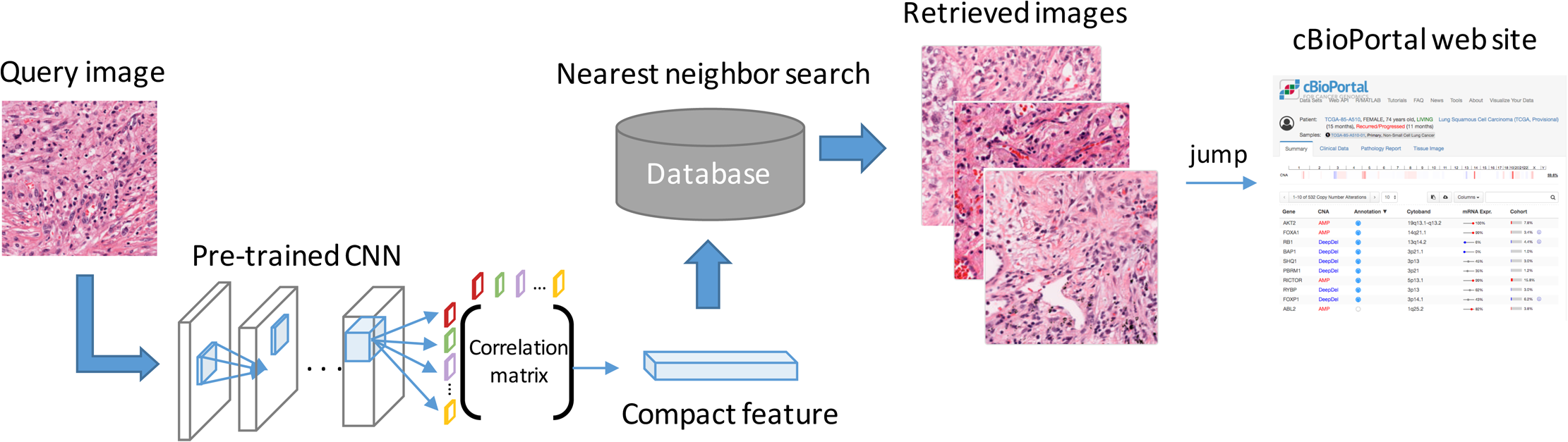
Overview of the Luigi system. After submitting the query image, deep texture representations are calculated and compressed by compact bilinear pooling, and the most similar images are retrieved. Users can view the retrieved cases on the cBioPortal website by clicking the link to the selected cases.

The Luigi system can be used by accessing its URL via any modern web browser. Users can search relevant images relevant to a query image from the database using intuitive operations. Users can also view the cBioPortal[11] website to view clinical, pathological, and molecular information relevant to the retrieved case. Although users do not need to register accounts to use the Luigi system, if an account is registered, the system saves a history of all uploaded images.

The Luigi database contains WSI information for each case; however, the query and retrieved images handled by the system are patches extracted from WSIs because in most situations, users submit small regions of a whole slide with specific features[4].

Details about the system are described in the following sections.

### Retrieval system

A CBIR system requires quantitative metrics that represent image “relevance.” Therefore, effective features must be extracted to compute the similarity between histopathological images. For example, the similarity of histopathological images is often considered within the context of cell structure, tissue texture, or shape of the nucleus. Many researchers have applied CBIR to medical domains by employing hand-crafted features or classical texture features based on such information. In contrast, the Luigi system uses deep texture representations computed using a pre-trained CNN for feature extraction of images, an approach motivated by recent studies of neural style transfer[9]. The deep texture representations are computed using a correlation matrix of feature maps in a CNN layer. Here the “conv3 1” layer of VGG-16[12] was employed.

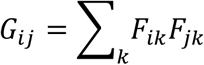

In the above equation, *G, F*, and *k* represent the correlation matrix, the feature maps of the CNN, and the position in the images, respectively. This manipulation, which is also called bilinear pooling, produces orderless image representations by aggregating the averages of the location-wise outer products of the feature maps. Note that this process discards some spatial information; therefore, the representations have spatial invariance, which appears to be necessary for pathological images because the similarity between two images should not be affected by the relative location of cells within the image. Because the representations are too high dimensional for practical application, the Compact Bilinear Pooling (CBP)[13] using Random Maclaurin approximation was adopted to approximate the original representations of 65536 dimensions with 1024 dimensions.

CBP is an approximation method of bilinear pooling using two random matrices and has good properties without the need for pre-training, unlike other dimension reduction methods, such as principal component analysis. Before the final representation, we added an element-wise signed square root layer and an instance-wise *l*_2_normalization as applied in the original paper [13]. The final representations for each image have 1024 dimensions, which demonstrated good performance when evaluating histopathological images.

When a query image is input into the Luigi system, deep texture representations of the original image, as well as the one rotated by 90° for imposing rotational invariance, are created and compressed using CBP. The system then searches for the nearest images in the database using the cosine similarity of the extracted compressed features between the query and database images.

To enable high-speed searching, an approximate nearest neighbor search method with the randomized kd-tree algorithm implemented in pyflann was adopted.

### Dataset and pre-processing

An image database was created from WSIs of all cancer types in the TCGA. The SVS files of the diagnostic WSIs that include DX in the file name were downloaded from the National Cancer Institute GDC legacy archive[14]. WSIs without sufficient information, such as magnification level, were discarded. At least three representative cancer regions for each slide were selected by professional pathologists using a Web browser-based software developed for this purpose (Figure 2). To avoid discordance between pathologists, annotations for each cancer type was performed only by a single pathologist. 342 slides were additionally removed due to poor staining, low resolution, out of focus across a slide, lack of cancerous regions, or likely to be mislabeled. Currently, the database contains WSIs of a total of 6763 patients in 32 cancer types (Table 1). Next, from each annotated region, 512 × 512 pixel patches at 40x magnification level was cropped from the regions and resized to 256 × 256 pixels. The position and the angle of the image patch was selected at random. Blur, scale and stain augmentation were applied and 60 image patches were generated from each patch to improve sensitivity (Figure 2). Finally, deep texture representations were calculated for the image patches and stored in the database.

**Figure 2.**
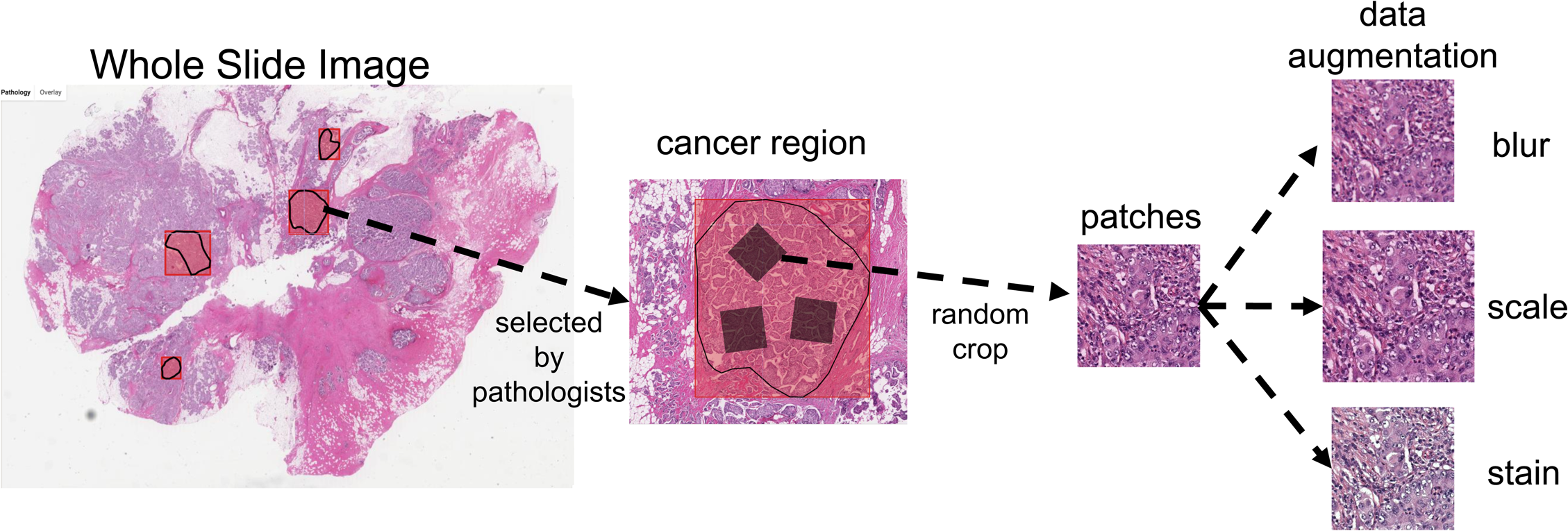
Pre-processing of the WSIs.

**Table 1.** Cancer types included in the current database. (placed at the end of this file)

| Cancer type | No. of WSIs |
| --- | --- |
| Breast invasive carcinoma | 811 |
| Glioblastoma multiforme | 513 |
| Lung squamous cell carcinoma | 502 |
| Lung adenocarcinoma | 473 |
| Brain Lower Grade Glioma | 461 |
| Sarcoma | 451 |
| Uterine Corpus Endometrial Carcinoma | 419 |
| Kidney renal clear cell carcinoma | 381 |
| Thyroid carcinoma | 381 |
| Head and neck squamous cell carcinoma | 341 |
| Colon adenocarcinoma | 339 |
| Bladder urothelial carcinoma | 337 |
| Prostate adenocarcinoma | 328 |
| Stomach adenocarcinoma | 291 |
| Liver hepatocellular carcinoma | 289 |
| Cervical squamous cell carcinoma and endocervical adenocarcinoma | 208 |
| Testicular germ cell tumors | 201 |
| Adrenocortical carcinoma | 166 |
| Pancreatic adenocarcinoma | 143 |
| Skin cutaneous melanoma | 127 |
| Ovarian serous cystadenocarcinoma | 84 |
| Kidney chromophobe | 82 |
| Mesothelioma | 69 |
| Pheochromocytoma and paraganglioma | 45 |
| Cholangiocarcinoma | 30 |
| Lymphoid neoplasm diffuse large B-cell lymphoma | 27 |

### User interface of the Luigi system

Figure 3 shows the interface of the Luigi system. Users upload a histopathological image file by dragging and dropping the image file, pasting a screenshot of the image, or clicking “click here to select a file”. The uploaded image is resized to 256 × 256 pixels, which is the same as the size of images in the database. Because this resizing process could change the texture, it is important to upload images with appropriate magnification levels to retrieve good results. Then the Luigi system will retrieve the most relevant patch in each WSI, rank each according to the cosine similarity, and initially display the relevant cancer cases among all cancer types. Tissues of origin in cancer can be changed by selecting the tissue from the pull down menu on top of the results. When a retrieved image is clicked, the selected image is set as a new query and can be further input to a query by clicking “Search” button. When the TCGA slide ID (TCGA-XX-XXXX-XXX-XX-XXX) below each of the image is clicked, a new window will open and the summary of the selected case will be displayed on the cBioPortal website, which enables the user to investigate the pathology report as well as clinical and molecular information of the case.

**Figure 3.**
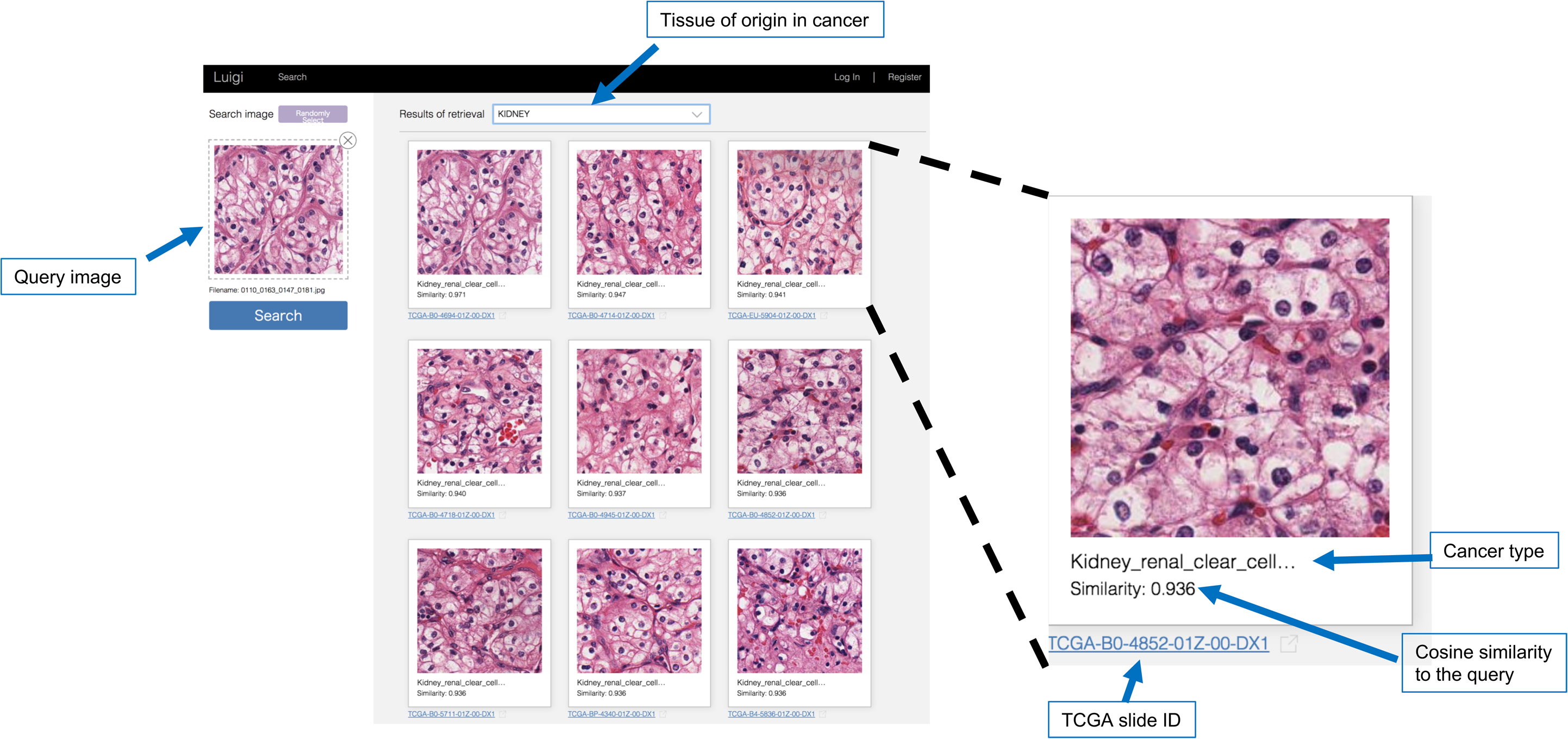
Graphical interface of the Luigi Webserver.

### Smartphone application

A smartphone application was also developed that allows users to directly submit appropriately scaled query images into the Luigi web application and view the retrieved images on a smartphone. Figure 4 shows the interface of the Luigi smartphone application. Users can take a photo of their own cases by tapping a button below on the screen or select a image stored in the smartphone. Users can easily adjust the size of a query image using above reference images. After touching a check button, the relevant cases of all tumors ordered according to the cosine similarity of the uploaded image will be displayed. Tissue of origin in cancer can be changed by clicking the above drop down menu on the screen. Users can also access the cBioPortal website by selecting a TCGA slide ID of the retrieved case. Currently, only an iOS version of the smartphone application is available, but an Android version will be developed in the future.

**Figure 4.**
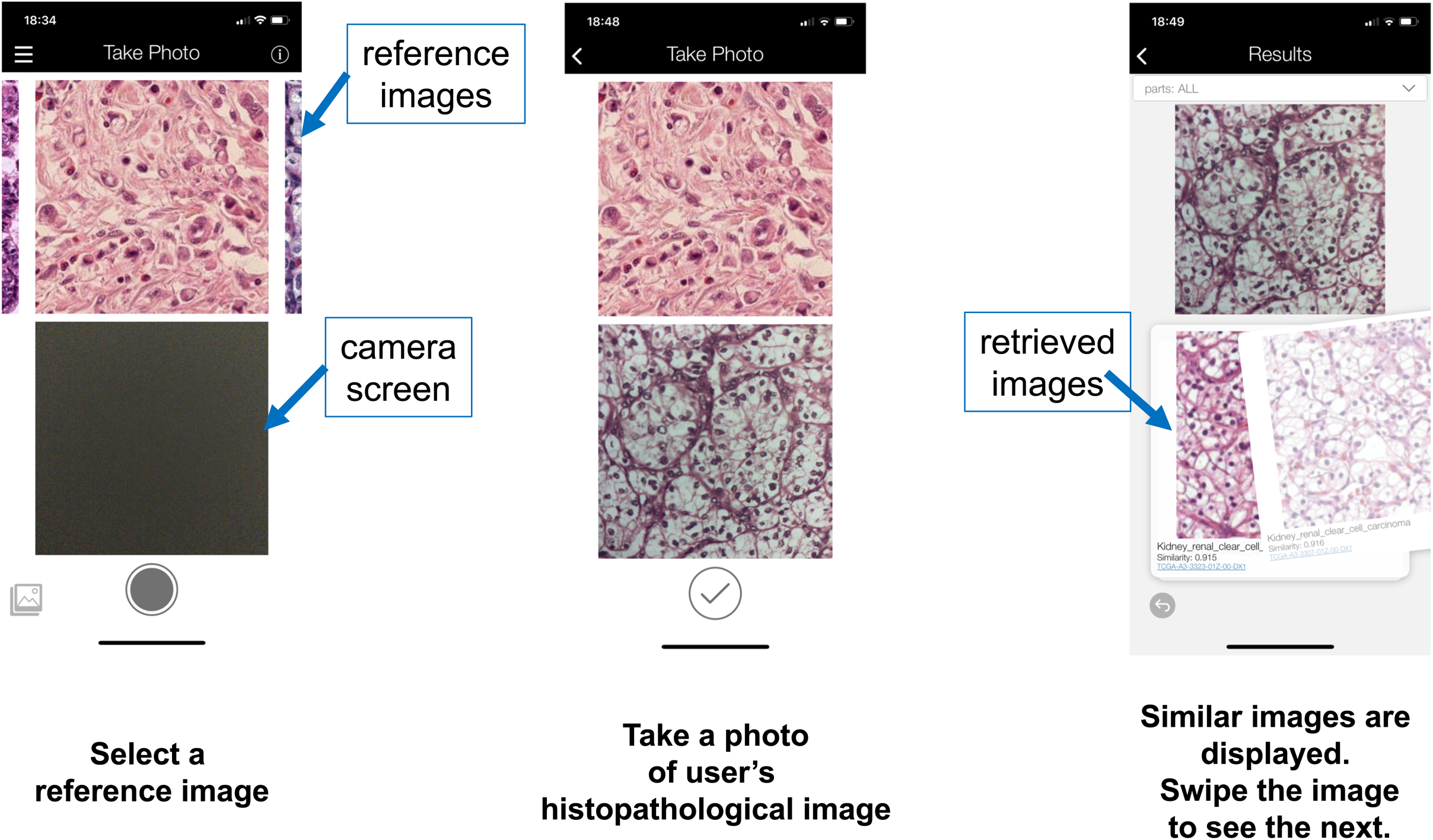
Graphical interface and usage of the Luigi smartphone application.

## Results

To determine if the Luigi system can accurately retrieve relevant images, the effectiveness of the Luigi system was assessed by querying some typical cancerous regions from TCGA images. Queries image used in this experiment are shown in Figure 5. For each query, the most relevant images in each WSI are retrieved and ranked according to the cosine similarity and then the averaged precision of the top 5, 10, and 20 images for three queries in each query cancer type is calculated to measure the retrieval accuracy. The query images, which show the typical appearance of each query cancer type, were selected at random and the retrieved images were evaluated based on its category or evaluated manually by two pathologists and a medical student.

**Figure 5.**
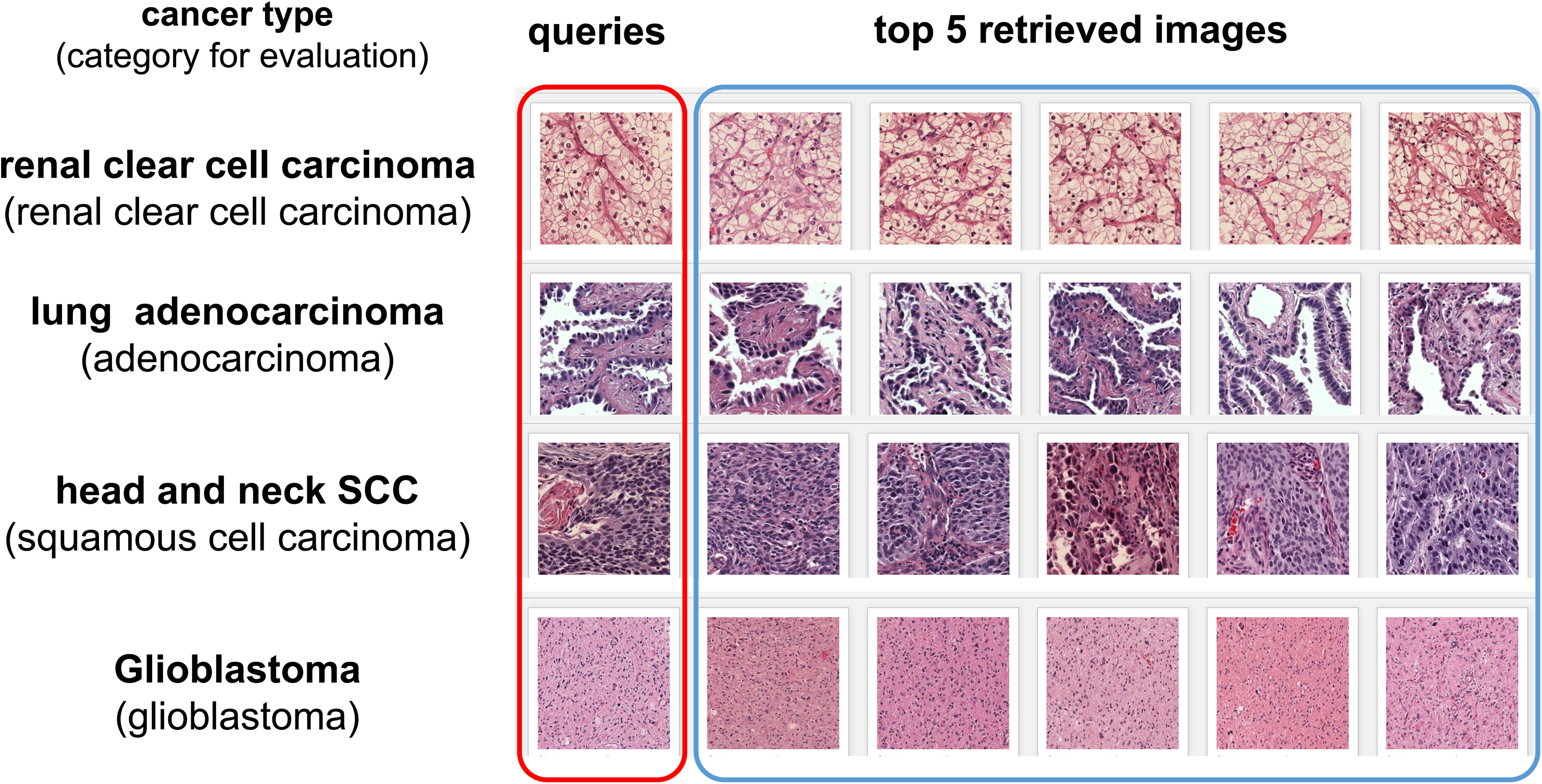
Query images and top 5 retrieval results of four cancer types used in the experiment.

Figure 5 and 6 present the top 5 retrieved images and the precision of the top 5, 10 and 20 retrieved images by the Luigi system, respecitvely. Most of the images appeared similar to the query image even to the pathologists, especially in top 10 retrieved images. The precision decreased as the number of retrieved images increased, but the precision keeps ≥ 50% even in top 20 images. These results demonstrate that the Luigi system successfully retrieved relevant images regardless of the cancer type, which is the purpose of the system.

**Figure 6.**
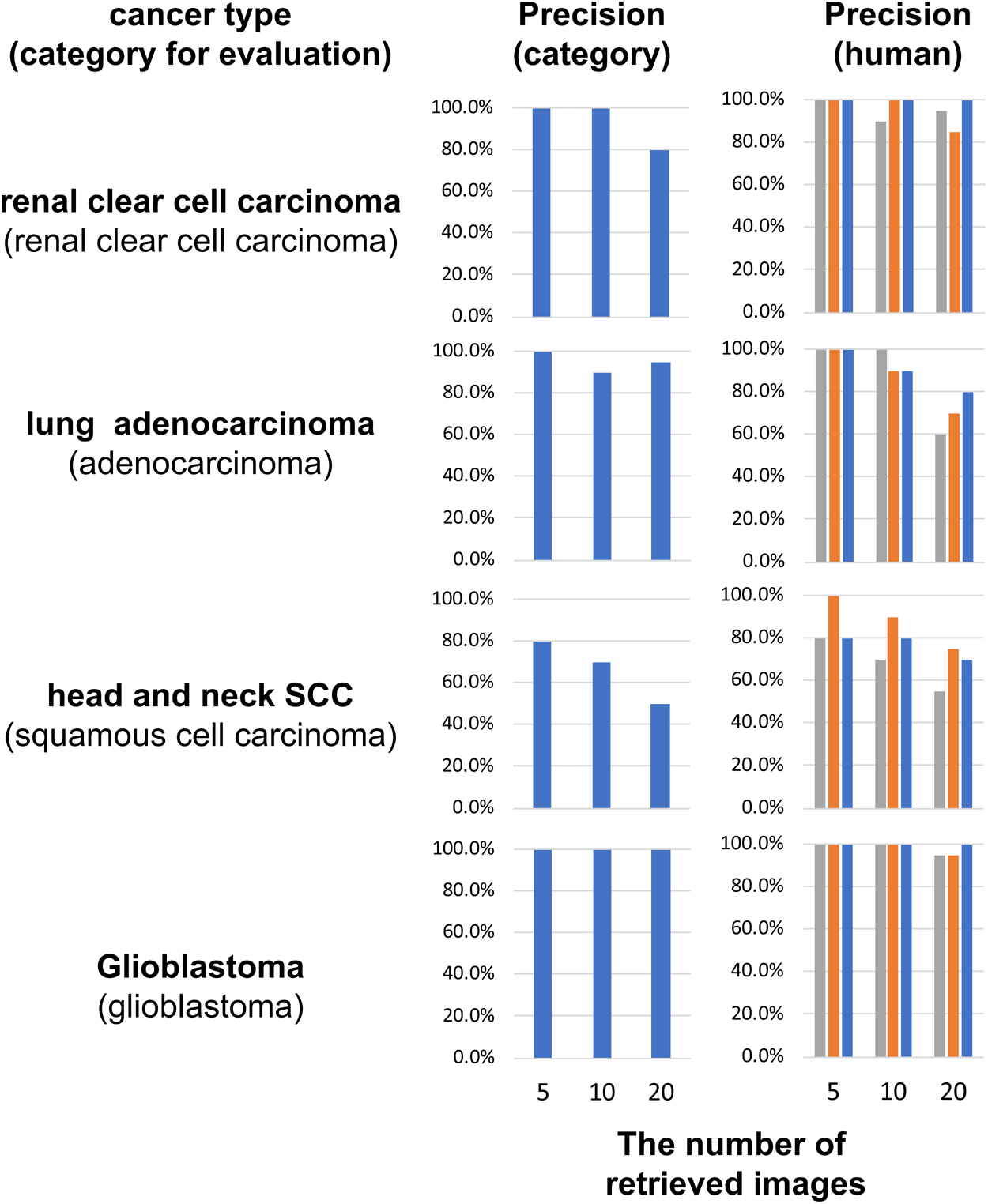
Precision of Luigi on The Cancer Genome Atlas dataset. Left: precision value based on the category. Right: precision value based on human evaluation (two pathologists and a medical student).

## Conclusions

We developed a novel web-based CBIR application, the Luigi system, which enables pathologists, students, and researchers to retrieve cancer cases with histopathological images similar to those of the query image. Experimental results showed that the Luigi system successfully retrieved relevant images. The features of Luigi system are as follows:

- To the best of our knowledge, this is the first publicly available CBIR system with rich clinical and genomic information in the pathology field.
- The Luigi image retrieval system uses deep texture representations to calculate the relevance metrics between pathological images.
- This enables the Luigi system to retrieve relevant images regardless of spatial information, e.g., the relative location of cells within the image.
- The Luigi system does not require extra learning; thus, it is scalable to a large number of histopathological images and any type of tissue or cell can be searched without training.
- The smartphone application allows users to easily submit query images without the need for special equipment and view the retrieved results on a smartphone.

We plan to evaluate the retrieval accuracy of the deep texture features from the CNN model, which is pre-trained in unsupervised way or fine-tuned using histopathological images. Each method has several issues to be resolved to improve the performance, but these are beyond the scope of this paper. For example, there are several options and hyperparameters for pre-training CNN using unsupervised learning such as learning methods (e.g. Restricted Boltzmann Machine, autoencoder, or Generative Adversarial Network), network structures, and learning rates. Fine-tuning could help improve the performance if histopathological images can be prepared with appropriate labels of various tissue types with an unbiased way. Although these ideas have not yet been implemented, we believe that the performance of the Luigi system is both satisfactory and practical.

## Availability and requirements

Project name: Luigi

Webserver: https://luigi-pathology.com/

iPhone application: available on the Apple iTunes store as “Luigi”

Operating systems: Platform independent (Webserver), iOS 10.0 or later (iPhone application)

Programming language: Python and Clojure (Webserver), Swift (iPhone application)

Other requirements: Firefox, Chrome, IE, or Safari. JavaScript must be enabled (Webserver).

License: None

Any restrictions to use by non-academics: Luigi cannot be used for commercial use.

## Abbreviations

CBIR, Content-based image retrieval; CNN, convolutional neural network; TCGA, The Cancer Genome Atlas; WSI, whole slide image; CBP, Compact Bilinear Pooling.

## Declarations

### Ethics approval

All image data related to human subjects used for this study is de-identified and publicly available from The Cancer Genome Atlas project. Therefore, this research is not classified as a human subject research and no Institutional Review Board approval is required.

### Consent for publication

Not applicable

### Availability of data and materials

The image datasets analyzed in this study are available in the National Cancer Institute GDC Data Portal, https://portal.gdc.cancer.gov/

### Authors’ contributions

D.K., K.F. contributed to the development of the web application; K.T. and D.K. contributed to the development of the smartphone software; A.K and H. Koda selected the cancerous regions in WSIs. R.S, T. U., H. Konishi, preprocess the image data. D.K. performed the experiments. T.H. and S.I. supervised the research; K.F. and D.K. led the writing of the manuscript with input from the other authors.

